# Dynamics of cell death due to N and P starvation across intra-clade diversity in *Prochlorococcus*

**DOI:** 10.1101/2025.05.24.655613

**Authors:** Yara Soussan-Farhat, Shira Givati, Osnat Weissberg, Waseem Bashir Valiya Kalladi, Dikla Aharonovich, Daniel Sher

## Abstract

Nutrient starvation and subsequent mortality are processes that can shape ecosystem dynamics and influence global biogeochemical cycles yet are poorly understood. Here, we examined the dynamics of culture decline in 15 strains of *Prochlorococcus*, globally abundant marine cyanobacteria, under nitrogen (N) and phosphate (P) starvation. We then ask whether mortality patterns can be related to the evolutionary history of each strain, the geographic location, and environmental conditions where it was isolated from, or the copy number of specific nutrient acquisition genes. We observed diverse decline patterns across starvation conditions and strains, identifying three differential features: maximum culture fluorescence, the number of fluorescence peaks during the decline stage, and the decline rate. Based on these features, each strain was categorized as more sensitive to either nitrogen starvation or nitrogen and phosphorus (mixed). High light (HL) strains are more sensitive to N starvation, whereas other facets of the strains’ evolutionary or ecological origin were not correlated with mortality features. The number of genes known to be involved in either N or P acquisition in each genome was not correlated with starvation sensitivity. Rather, genes involved in DNA damage repair were associated with N sensitivity to starvation, especially in HL strains, whereas genes related to protein quality control were more abundant in LL strains and associated with mixed sensitivity. These findings reveal a previously unrecognized diversity in the dynamics of starvation and mortality across closely related *Prochlorococcus* strains, potentially driven by differences in the responses to DNA and protein damage.

## Introduction

Death of microbes might be due to ecological processes, such as predation and phage infection, or they can die from “natural” causes, including oxidative stress, elevated temperature, and nutrient depletion [1–3]. Although death due to predation or phage infection has been intensively studied (e.g. [1–3]) much less is known about natural mortality, yet the rate at which bacterial cells die, and potentially how they die, can have important biogeochemical consequences. This is because the release of cellular contents from dying cells provides organic matter that can be recycled within microbial communities (“the microbial loop”; for examples from the marine environments see [4–6]). In aquatic systems, cells that die but are neither internalized (eaten) by other organisms nor immediately lysed may also influence aggregation and vertical flux, contributing to the export of organic carbon and nutrients to deeper waters (e.g. [7,8]). These fluxes are central to the regulation of atmospheric CO₂ and the transfer of bioavailable material to deeper ecosystems [9].

One potential cause of microbial death is starvation [10,11]. Different microbes respond differently to starvation, for example, in *E. coli*, slower growth under carbon starvation enhances survival [12], whereas *B. subtilis* forms stress-resistant spores under nutrient limitation [13]. Different stresses also interact, leading to trade-offs; for instance, *E. coli* cells selected for UV resistance also show increased starvation tolerance but become more vulnerable to salinity changes [14]. Despite these observations, natural mortality patterns in common environmental microbes, particularly marine bacteria, remain poorly characterized. In particular, how these mortality dynamics vary across changing ecological and evolutionary contexts represents a key knowledge gap.

Here, we characterize the dynamics of starvation and mortality in *Prochlorococcus*, a globally abundant marine cyanobacterial clade, which is the numerically dominant primary producer in oligotrophic regions [15]. *Prochlorococcus* has the smallest genomes of all primary producers, providing a model for the simplest biological system capable of living off sunlight and inorganic nutrients [15–17]. The *Prochlorococcus* “federation” comprises many diverse lineages, each equipped with its unique set of genes allowing it to thrive in its oceanic niche [15]. These lineages are typically divided into two main categories: high-light-adapted strains (HL), that grow at the ocean surface, where nutrients are scarce, and low-light-adapted strains (LL) found in deeper waters where nutrients are more abundant. All *Prochlorococcus* strains (HL and LL) share ∼1,000 core genes for essential housekeeping functions [18], with thousands of additional “flexible” genes driving lineage-specific adaptations, giving rise to at least six ecotypes (HLI–HLVI, LLI–LLVII) [15,18,19]. Differences in responses to environmental stresses, such as nutrient stress or co-stresses, can be understood, at least in part, through this core-versus-flexible gene content. Specifically, about 92 genes have been shown to be involved in the response of several *Prochlorococcus* strains to N and P limitation or starvation and these genes have been used as biomarkers to infer nutrient limitations in the ocean [20–23].

Despite the work and characterization of the short-term response to starvation in *Prochlorococcus*, little is known about the subsequent, mostly unexplored stages, of long-term starvation. Previous studies have shown that carbon- and nitrogen-starved cells die rapidly, and that the rate of mortality is reduced when cells interact with some heterotrophic bacteria, but not others [24–27]. However, to what extent the dynamics of starvation and mortality differ between different stressors, and whether they are related to the evolutionary and ecological history of different strains, remain unknown. We posit that a better understanding of starvation and mortality in *Prochlorococcus* could help understand basic aspects of cell biology and physiology in a “simple” model bacterium and could also provide constraints on the rates of mortality under natural conditions. Indeed, the fraction of *Prochlorococcus* cells in the ocean dying from “natural” sources (i.e. neither phage predation nor grazing) is unknown, but estimates range from less than 1% to more than half (e.g. [28,29]). To begin addressing these questions, we cultured fifteen axenic *Prochlorococcus* strains, isolated from different oceanic regions and representing broad genetic diversity, under conditions where cessation of growth and subsequent mortality are due to N and P starvation. We show that these strains exhibit diverse “death phenotypes” which can be partly quantified using specific features of the growth-and-death curves-the maximum culture fluorescence, the number of fluorescence peaks during the decline stage, and the decline rate. We propose that these features are associated primarily with the HL-LL division and less with the specific ecological and evolutionary history of each strain. These features were also not associated with known N and P acquisition genes, but rather with genes associated with DNA and protein quality control and repair.

## Material and methods

### Experimental design

Fifteen axenic strains of *Prochlorococcus* were maintained under constant light (22-25 μmole photons m^−2^ s^−1^) at a temperature of 22°C. Four distinct nutrient conditions were prepared from natural seawater collected at a depth of 20 meters, using replete media (PRO99) as a baseline [30], alongside three nutrient-depleted variations: a low nitrogen medium (lowN, reduced 8-fold) and two distinct low phosphate media (lowP, reduced 8-fold and 50-fold, respectively). For nutrient concentration please see Supplementary Table S1.

Due to the large number of strains, conditions and replicates, the experiment was divided into four distinct sub-experiments, each consisting of 3-4 different strains. *Prochlorococcus* strains MED4 and MIT9312 were used as internal standards across all sets (and therefore these strains have 12 replicates for each media type). The experiment unfolded in two stages. Stage 1 (the conditioning stage) spanned one week that served as a nutrient-depletion phase; cultures were transferred once from replete media directly into their respective nutrient-limited media to deplete internal cellular stores and minimize nutrient carry-over. All other conditions were identical. The axenic status of the cultures was also verified alongside stage 1 by introducing 1 mL of culture into 15 mL of PROMM medium [30] and incubating for 7 days. These test tubes were maintained alongside the conditioning phase of the experiment under identical constant light and temperature conditions. Cultures were examined visually for turbidity; any strain exhibiting visible growth was classified as non-axenic and excluded from the experimental phase (stage 2). In total, two strains which were initially thought to be axenic (MIT9303 and MIT0609) were identified as contaminated and were excluded from the experiment. Prior to initiating Stage 2 (the experimental phase), cell abundance was quantified using flow cytometry, and the initial concentration for all experimental cultures was set to 2 × 10^6^ *Prochlorococcus* cells per mL. Each strain-by-medium combination was established in triplicate.

Daily monitoring of the strains’ growth was conducted by measuring fluorescence (FL-ex440; em680) using a Fluorescence Spectrophotometer (Cary Eclipse, Varian) as a proxy for cell density. Cultures were sustained until all reached a fluorescence level of less than 0.1, this threshold was chosen as a conservative limit of detection, remaining well above our measured baseline background (mean blank fluorescence = 0.013; *n* ≈ 150), with the experimental duration presented in this study capped at 90 days post-setup.

### Bulk fluorescence as a proxy and assessment of culture axenicity

Two separate experiments were conducted to determine whether measuring bulk fluorescence (FL) serves as a reliable proxy for cell abundance, given that physiological factors such as chlorosis and shifts in cellular fluorescence per cell can alter bulk FL measurements. Four to five representative strains were cultured across the four tested nutrient conditions (detailed in Supplementary Text S1 and Supplementary Text S2). Samples for flow cytometry were collected at the initial start point, peak growth, and the subsequent decline phase. Samples were fixed with glutaraldehyde (0.125% final concentration), incubated in the dark for 10 min, and stored at −80 °C until analysis. Fluorescent beads (2 µm diameter, Polysciences, Warminster, PA, USA) were added as an internal standard, and data were acquired and processed using FlowJo software.

Axenicity tests were performed using PROMM and MB media; tubes were incubated under identical conditions to the experimental setup, and cultures were visually examined for turbidity. Any strain exhibiting visible growth was classified as non-axenic and excluded from the analysis. Additionally, we sequenced 16S rRNA amplicons from the experimental strains prior to setup and on randomly selected samples at the end of the second experiment (Supplementary Text S3). Because these control experiments resulted in patterns similar to the original experiment (Supplementary Figure S3) and were axenic, we conclude that the patterns observed in the original experiments were likewise unaffected by contamination.

### Calculating specific growth and death characteristics

#### Growth rates

Growth rates (μ) were determined by fitting a linear regression to the log-transformed exponential phase data, expressed as ln(N_t_) = ln(N_0_) + μ(t − L). In this model, N_t_ is the cell concentration at time t, N_0_ is the initial concentration, and L is the lag phase duration. Growth rates were accepted only for models with a coefficient of determination (*R*^2^) > 0.9, where μ was defined as the regression slope [25].

#### Normalized FLmax

To enable cross-comparison between distinct clades and ecotypes, maximum fluorescence (MaxFL) values were normalized against a reference condition. We calculated a baseline for each strain by averaging the MaxFL to that of the replete conditions (PRO99). Each individual MaxFL value was then expressed as a percentage of this group-specific mean:

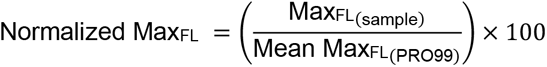

The standardization accounts for inherent variations in maximum yield across different lineages, allowing for a relative assessment of growth performance under experimental conditions.

#### Peak count

The peak count was determined by applying the ‘find_peaks’ function from the SciPy package in python (1.15.0) with specified parameters: height=0.15 and threshold=0.05. A total of 18 distinct parameter combinations were systematically tested, including an additional parameter-distance, which was set to none. The optimal parameter set, yielding minimal noise, was selected. The ‘max’ function, a built-in component, was integrated alongside the ‘find_peaks’ function to guarantee the identification of the highest peak in all instances.

#### D-value

The D-value is defined as the time required to reduce cell population by 90% [31]. Here, the systematic error (0.15) was subtracted from the FLmax to determine the D-values from the maximum growth. Tmax is defined as the day the cultures reached their maximum yield, T10% is the time taken to reach 90% decline.

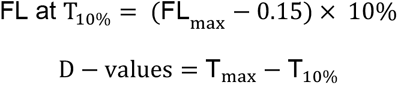

To test how growth rate, MaxFL, peak count, and D-values, differ within each strain across various media, we employed Ordinary Least Squares (OLS) regression using the statsmodels package (python v0.14.4). For each strain, we modeled the dependent variables as a function of the growth media. To control for the family-wise error rate associated with multiple testing, pairwise comparisons were conducted using the Benjamini-Hochberg procedure to adjust *P* values.

### Testing correlation between environmental conditions, N/P acquisition genes, COGs, and starvation sensitivity

#### Environmental conditions

We performed an extensive literature search to obtain the environmental conditions measured when each *Prochlorococcus* strain was isolated. When such data were unavailable, we contacted the authors. Nevertheless, data for certain strains were unavailable, therefore the World Oceanic Atlas [32], which provides interpolated data, was used for obtaining phosphate, nitrate, and temperature measurements specific to the isolation origin and timing for each strain (Table 2, Supplementary Figure S8).

Data were collected from monthly records, selecting the closest location to the actual isolation site. Measurements were taken within a 10-meter range on either side of the isolation depth (e.g., if a strain was isolated at 15 meters, data were collected from 5–25 meters). The average was then calculated for this range. No significant difference was observed between the data from the WOA and those available during the strain isolation (Supplementary Figure S8).

#### N and P acquisition genes

To test whether starvation sensitivity was related to the genetic capacity to take up N or P resources, we selected 50 N-related genes associated with the utilization of ammonium, nitrate, nitrite, urea, amino acid and molybdopterin biosynthesis [22,33,34]. Additionally, 47 P-acquisition genes were chosen for their role in phosphate, phosphite, phosphonate and phosphoanhydride metabolism [20,35–39]. The presence and frequency (measured by copy number) of these genes was assessed in each strain using the phyletic pattern data in the Cyanorak [40] (Dataset 2). A test of these genes across multiple published transcriptome datasets of N and P starvation experiments confirmed that the expression of these genes indeed changed under the appropriate starvation conditions (see “integrated transcriptome analysis”, below, Supplementary Text S6 and Supplementary Figure S9A).

#### COGs across genomes

To obtain COGs data, the CyanoRak database was accessed, which provides GFF files describing the genomes of the selected *Prochlorococcus* strains, apart from MIT1314 and MIT1327. COG numbers were extracted from these files using a Python script (3.12.8).

MIT1327 was not fully annotated; however, COGs were retrieved from its GF file provided by the CyanoRak team. MIT1314 was excluded from this analysis as it has not yet been annotated.

To identify significant differences in gene copy numbers and environmental measurements between distinct strain groups (such as HL/LL or N/mixed sensitive), we employed the Mann-Whitney U test. The non-parametric approach was selected to compare the underlying distributions of the two groups, making it robust to potential outliers and non-normal data distributions. For each measurement, we calculated the median values for both groups to determine the directionality of the observed effects. To account for the inflation of Type I error resulting from multiple comparisons, all *P* values were adjusted using the Benjamini-Hochberg (FDR) procedure.

### Integrated analysis of multiple transcriptome studies

A literature search was performed using a bespoke knowledge graph (KG) integrating data extracted from approximately 40 published studies on *Prochlorococcus* and related organisms, with each gene also annotated using Cyanorak, eggnog-mapper, and RefSeq. The KG is integrated with Claude Code via the Model Context Protocol (MCP), constraining the model to rely exclusively on the curated KG rather than internal memory and backing all downstream analyses with reproducible Python scripts. A full description of the KG, including code, is available on Github (https://github.com/TheSherLab/15strain4media_analysis), and the detailed analysis is described in the Supplementary Text S6. A publication describing the KG is in prep.

## Results and discussion

### Diverse growth and death patterns across strains and starvation types

*Prochlorococcus* is usually cultured in media where the main macronutrients N and P are balanced, i.e. provided at the 16:1 ratio found in average biomass across the oceans (the Redfield Ratio, [41]). Under such conditions the limiting factor leading to cessation of growth is unknown and can include carbon limitation and/or toxicity due to metabolic byproducts ( [30,42], Supplementary Text S2 and Supplementary Figure S2). Therefore, to measure how N or P starvation affect *Prochlorococcus* growth and death, we modified the commonly used PRO99 medium by reducing NH4^+^ concentrations 8-fold (“lowN”, [43]) or PO4^3-^ concentrations (8- or 50-fold “lowP”, [44]). We chose two different concentrations of PO4^3-^ because *Prochlorococcus* are known to significantly reduce their P requirements (e.g. through the production of sulfolipids [45]), leading to highly variable N:P ratios [46], and we hypothesized that an 8-fold reduction might not induce P starvation in all strains. These media were used to culture 15 axenic *Prochlorococcus* strains, which were isolated from different oceanic locations and represent much of the genetic diversity of this globally abundant clade (Figure 1A, B and Table 1).

**Figure 1:**
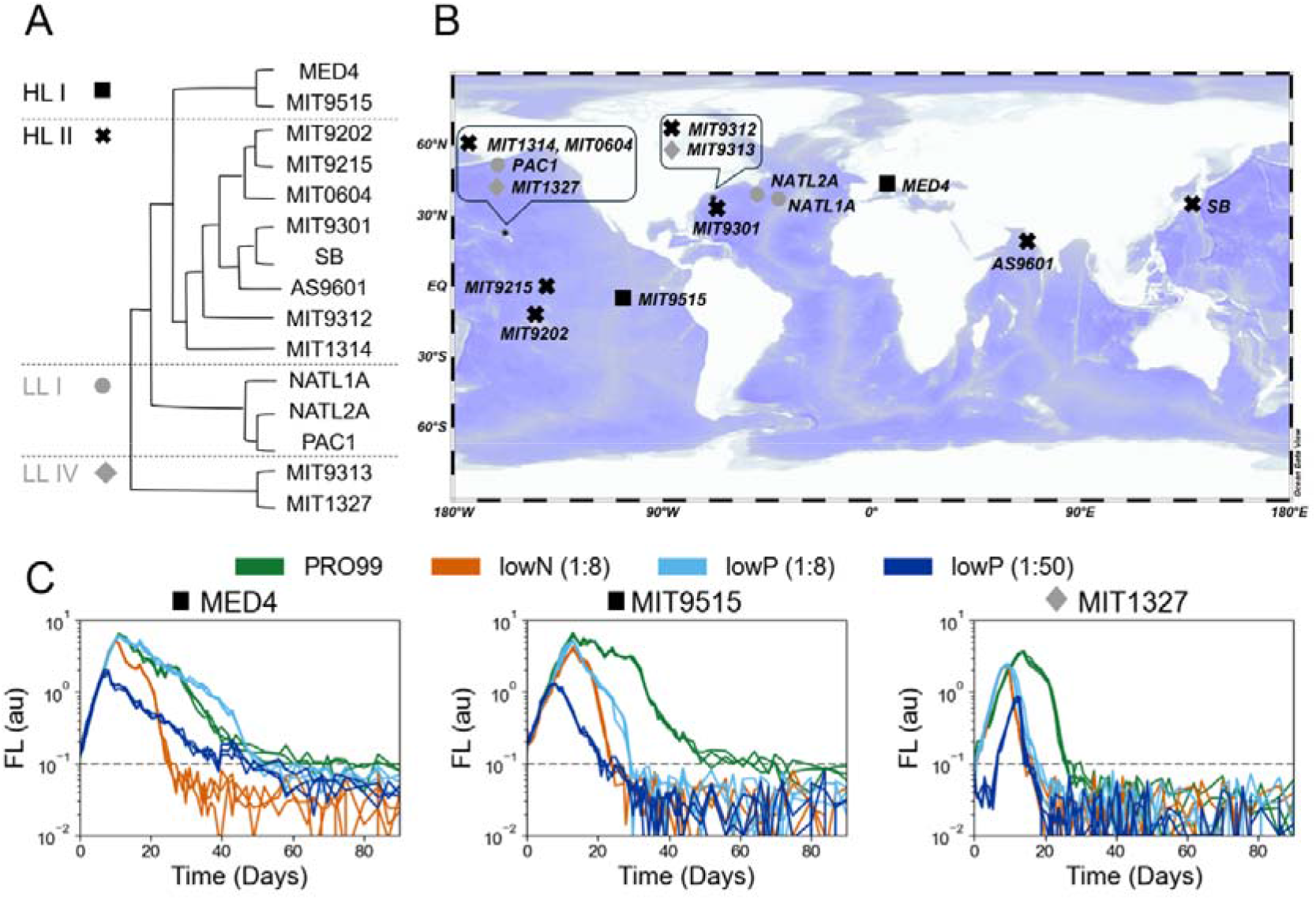
Characteristics of the 15 strains used, and examples of growth-and-death curves under multiple nutrient conditions. A. Schematic phylogenomic tree (based on [50,74]) illustrating the evolutionary diversity of the 15 strains used; black and grey represent HL and LL lineages, respectively, whereas shapes indicate distinct ecotypes. Dotted lines highlight the ecotype separation. B. A world map highlighting the isolation location of each strain. Multiple strains may have been isolated from the same location. C. Growth-and-death curves for three selected strains (see Supplementary Figure S4 for all curves); the x-axis represents time in days, and the (logarithmic) y-axis represents bulk culture chlorophyll autofluorescence (FL, arbitrary units). Individual lines represent biological triplicates. Colors represent the media under which cultures grew, and shapes above each plot denote ecotypes as shown in panel A. Highly variable fluorescence under 10^-1^ au is considered instrument noise and is marked with a dotted line.

**Table 1:** The axenic *Prochlorococcus* strains used in this study, with information on when and where they were isolated, environmental conditions at the time of isolation (from the World Oceanic Atlas), and sum of known N and P acquisition genes.

| Nitrate (μmol/kg) | Phosphate (μmol/kg) | P acquisition genes | N acquisition genes | Strain reference |
| --- | --- | --- | --- | --- |
| 2.36 | 0.11 | 34 | 22 | [47] |
| 2.84 | 0.40 | 14 | 11 | [48] |
| 0.78 | 0.05 | 17 | 18 | [49] |
| 5.73 | 0.76 | 16 | 19 | [49] |
| 2.85 | 0.18 | 20 | 39 | [50] |
| 0.36 | 0.03 | 37 | 19 | [48] |
| 2.15 | 0.17 | 24 | 32 | [51] |
| 2.90 | 0.63 | 16 | 19 | [52] |
| 1.47 | 0.36 | 27 | 22 | [53] |
| 0.93 | 0.13 | 20 | 18 | [54] |
| 0.87 | 0.13 | 28 | 25 | [19] |
| 3.17 | 0.11 | 28 | 25 | [55] |
| 0.08 | 0.08 | 19 | 31 | [56] |
| 0.93 | 0.18 | 22 | 25 | [53] |
| 1.47 | 0.36 | 17 | 23 | [57] |

| Strain | Clade | Isolation date | Isolation location | Coordinates | Depth (m) | Genome size | GC% | Temperature (°C) |
| --- | --- | --- | --- | --- | --- | --- | --- | --- |
| **MED4** | HLI | January 1989 | Mediterranean Sea | 43°N 6°E | 5 | 1,657,991 | 30.8 | 13.34 |
| **MIT9515** | HLI | June 1995 | Tropical Pacific | 5.7°S 107.1°W | 15 | 1,704,176 | 30.8 | 26.81 |
| **MIT9202** | HLII | September 1992 | Tropical Pacific | 12°S 145.4°W | 79 | 1,691,453 | 31.1 | 26.77 |
| **MIT9215** | HLII | October 1992 | Equatorial Pacific | 0° 140°W | 5 | 1,738,790 | 31.1 | 25.61 |
| **MIT0604** | HLII | May 2006 | North Pacific | 22.75°N 158°W | 175 | 1,780,061 | 31.2 | 20.04 |
| **MIT9301** | HLII | July 1993 | Sargasso Sea | 34.8°N 66.2°W | 90 | 1,641,879 | 31.3 | 20.05 |
| **SB** | HLII | October 1992 | Western Pacific | 35°N 138.3°E | 40 | 1,669,973 | 31.5 | 23.37 |
| **AS9601** | HLII | November 1995 | Arabian Sea | 19°N 67.10°E | 50 | 1,669,886 | 31.3 | 26.52 |
| **MIT9312** | HLII | July 1993 | Gulf Stream | 37.5°N 68.2°W | 135 | 1,709,204 | 31.2 | 19.54 |
| **MIT1314** | HLII | June 2013 | North Pacific | 22.75°N 158°W | 150 | 1,704,447 | 31.2 | 20.34 |
| **NATL1A** | LLI | April 1990 | North Atlantic | 37.4°N 40°W | 30 | 1,864,731 | 35.0 | 16.89 |
| **NATL2A** | LLI | April 1990 | North Atlantic | 39°N 49.3°W | 10 | 1,842,899 | 35.1 | 17.34 |
| **PAC1** | LLI | April 1992 | North Pacific | 22.75°N 158°W | 100 | 1,842,113 | 35.1 | 21.85 |
| **MIT9313** | LLIV | July 1993 | Gulf Stream | 37.5°N 68.2°W | 135 | 2,410,874 | 50.7 | 20.34 |
| **MIT1327** | LLIV | June 2013 | North Pacific | 22.75°N 158°W | 150 | 2,587,389 | 50.3 | 19.54 |

To ensure the cultures were indeed limited by the nutrient (or starved of the nutrient), we performed a spiking experiment in which cells were grown under the same conditions mentioned above but were spiked with the limiting nutrient upon reaching their peak. Spiking the cultures resulted in renewed growth comparable to that observed in PRO99 medium conforming that indeed growth stopped due to these nutrients becoming limiting (Supplementary Text S2 and Supplementary Figure S2).

The growth and death of each strain were followed over 90 days by measuring bulk culture fluorescence, a proxy for cell growth resulting in a dataset of 252 growth curves. The growth-and-death curves were highly reproducible both between biological triplicates (Figure 1C, Supplementary Figure S4) and between independent experiments with two strains (MED4 and MIT9312, Supplementary Figure S5 and Supplementary Text S4). Clear differences in the shape of the growth-and-death curves under N and P starvation, as well as between nutrient starved conditions and the PRO99 medium, were observed both within and between strains (Figure 1C, Supplementary Figure S4). Most strains showed no significant differences in exponential growth rates under lowN (1:8) and lowP (1:8) conditions, with similar growth observed in 80% of cases. However, under more severe P starvation (1:50), most strains exhibited a reduced growth rate (Supplementary Figure S6). Nevertheless, the main difference was visually seen in the mortality stage between the different nutrient starvation types, and the different strains, consistent with previous studies [25].

### Quantitative features of culture decline

Although the growth phase of most cultures could be described by a single, exponential growth rate, the stationary and decline phases are much more complex, with multiple peaks, troughs and decline stages (Figure 1C, Supplementary Figure S4). We therefore defined three quantitative features, each describing different aspects of the transition to nutrient starvation and subsequent mortality: the maximum fluorescence, number of fluorescence peaks, and the time it took the cultures to decline to 10% of maximum fluorescence (decimal reduction time, D-value, [31]). Each feature captures a distinct aspect of cellular death that is differently affected by starvation conditions in individual strains and between strains, (for statistical results see Supplementary Table S2 and for measurements see Dataset 1).

#### Maximum fluorescence yield-MaxFL

Bulk culture chlorophyll Fluorescence (FL) is an easily measurable, non-invasive parameter that, generally, is correlated with cell abundance during growth and decline stages (Supplementary Text S1 and Supplementary Figure S1, [25,27]). However, interpreting fluorescence maxima as direct measures of cell density requires caution, as chlorotic cells emerge under starvation. These cells are characterized by low chlorophyll autofluorescence and can exhibit reduced FL without necessarily indicating a decrease in cell abundance [24]. Nevertheless, a separate experiment demonstrated that differences in MaxFL within individual strains are primarily driven by absolute cell abundance rather than chlorosis, confirming that changes in peak fluorescence closely mirror true cell numbers (Supplementary Text S1, Supplementary Figure S1). Our findings indicate that MaxFL is strongly influenced by growth conditions, with the replete medium consistently producing the highest yield across nearly all strains. In 40% of cases, growth in lowP (1:8) resulted in a MaxFL comparable to PRO99. This suggests that an 8-fold reduction in P may not be sufficient to induce P starvation in these strains, with cessation of growth potentially due to other factors such as the accumulation of waste products (Supplementary Text S2 and Supplementary Figure S2). Additionally, even though lowP (1:50) reduced MaxFL compared to lowP (1:8), the decrease was not as large as the ∼ six-fold reduction expected from a simple linear relationship between available PO₄³⁻ and biomass. In contrast, MaxFL is lower in all strains grown in lowN than in PRO99, although the reduction was also generally less than the expected eight-fold decrease compared to PRO99 (MaxFL Figure 2B, see also [43]). Taken together, these results suggest that lowN (1:8) and lowP (1:50) induced N and P starvation, respectively, but that quantitatively assessing the impact of starvation on yield based on bulk culture autofluorescence may be complicated by varying physiological responses (e.g., chlorosis and, in the case of PRO99, additional mortality factors such as the exudation of toxic byproducts, [43]).

**Figure 2:**
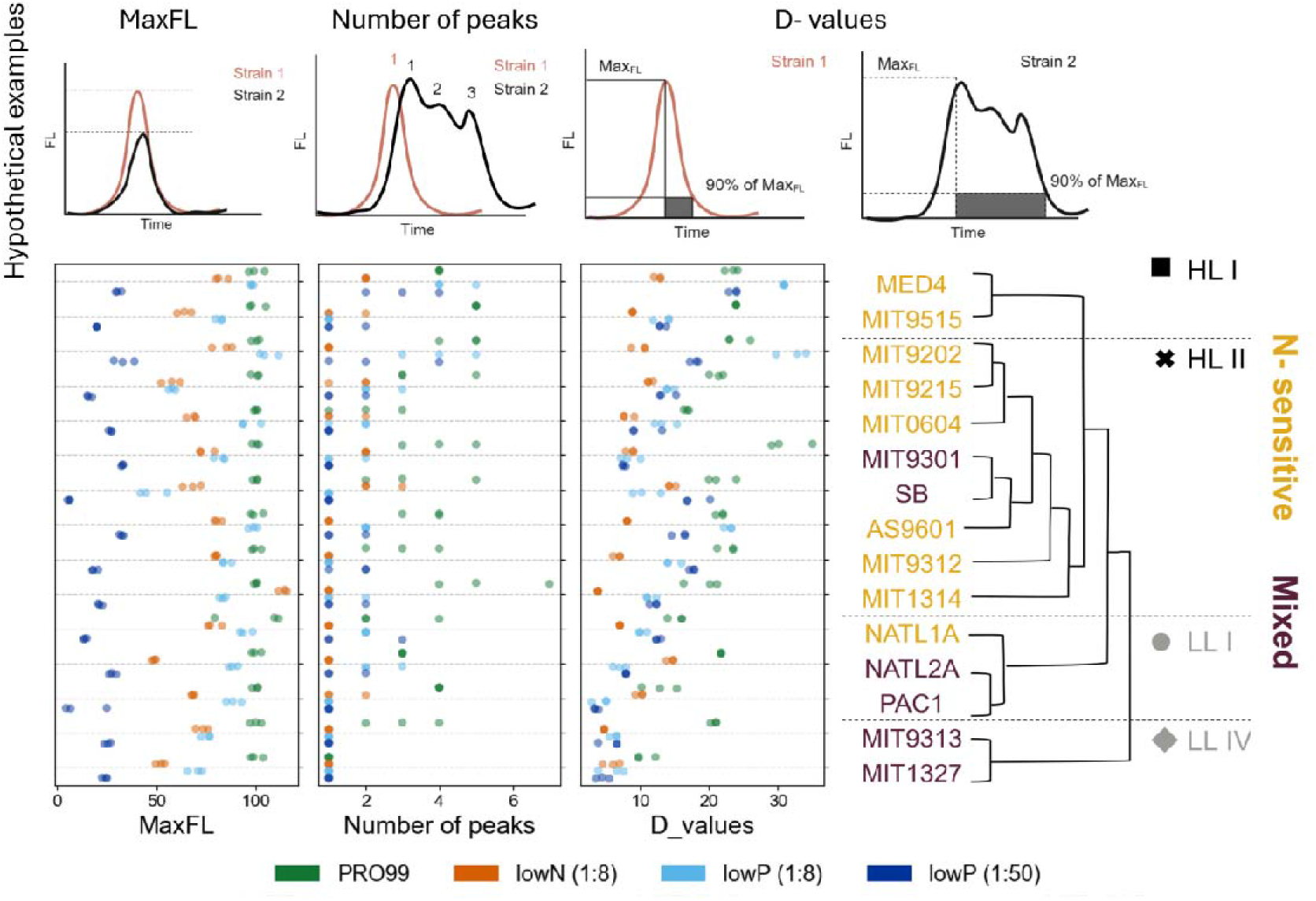
Features describing the culture decline stage. A. MaxFL; B. Peak count; C. D-values. The upper illustration shows a hypothetical difference between two stains as an example of the three features. Each circle represents a biological triplicate. Jitter was introduced to improve the visualization of overlapping points. Strain names are colored according to their sensitivity type, which is determined by the D-value (see text). A lower D-value indicates a faster decline in a given condition, reflecting greater sensitivity.

#### Peak count

As the population declines, multiple peaks and troughs are observed in the growth-and-death curves, and these patterns are reproducible across replicates and conditions (Figure 1, Figure 2B, Supplementary Figure S4). As noted above, these peaks may reflect changes in physiological properties such as fluorescence per cell, or phases where cell abundance actually increases. In *E. coli*, peaks and troughs in the growth and death curves can be due to the cells beginning to catabolize amino acids, carbohydrates, lipids, and even DNA from dying cells [58,59]. This process continues until all viable cells are exhausted, and (primarily at later stages, after most cells have died) may be accompanied by the emergence of genetically distinct lineages with mutations conferring a growth advantage under stationary phase (the “GASP” phenotype, [58]). In *Prochlorococcus*, we calculated the number of peaks from the point of MaxFL and onwards, observing that most of the peaks occur in the stationary or early decline phases, and mostly in the PRO99 media (Figure 2, Supplementary Figure S4). In another experiment where cell counts were measured, some cases were observed where fluctuations in FL corresponded to changes in cell number, representing true population re-growth rather than shifting cellular fluorescence (Supplementary Text S1, Supplementary Figure S1F). Given that these peaks occurred relatively early and when FL was high, it is less likely that they represent a true GASP phenotype. In contrast to most strains, three strains (PAC1, MIT9313 and MIT1327) did not exhibit multiple peaks under nutrient-deplete condition, and in MIT1327 they were not observed even in PRO99 (Figure 2B). What drives these changes in culture fluorescence, and what molecular processes define them; requires further investigation.

#### D-value

The D-value, a widely used metric in microbiology, refers to the time required for 90% of the cells in a culture to die [31]. A large D-value therefore reflects a slower decline rate and prolonged survival. Differences in the D-value were apparent across both treatment types within an individual strains and across strains. The PRO99 replete medium expectedly exhibited the highest D-value (longest survival), individual strain displayed contrasting mortality rates under N versus P starvation. Despite the low maximum fluorescence (MaxFL) and reduced yield under lowP (1:50), N-deplete conditions induced the fastest decline in 8 of the 15 strains. It is tempting to speculate that this reflects nutrient-specific adaptive strategies (e.g., sacrificing yield to prolong survival under P-stress versus experiencing rapid, irreversible decline under N-stress). A positive correlation (*r* = 0.71) was observed between the peak count and D-value, suggesting that dynamic changes during stationary stage (i.e. changes in per-cell fluorescence or in cell numbers, whether due to nutrient starvation or another stressor) may play a significant role also in determining death rate (Supplementary Figure S7).

### Sensitivity to starvation is associated with the HL-LL division in Prochlorococcus

We next asked whether the different patterns of mortality can be related to the evolutionary or ecological background of the different *Prochlorococcus* strains. The phylogeny of the strains (Figure 1A), and particularly the differentiation between HL and LL adapted clades, is associated with evolutionary differences in many traits such as genome size, metabolic complexity and, potentially, the capacity to utilize diverse organic compounds (Table 2, [15]). LL adapted strains have larger genomes and a higher predicted capability to utilize various organic resources such as amino acids and nucleotides and thus grow mixotrophically (using organic resources rather than only inorganic ones, [60–62]). We therefore expected these strains to be better at utilizing organic matter released from dying cells (“necromass”, [3]), resulting in slower decline (higher D-value), and potentially more peaks of “re-growth”. However, while the most apparent difference in D-values was indeed between the HL and LL adapted clades, it was the HL adapted strains that declined more slowly under P starvation (higher D-values, Figure 3A), suggesting they are more adapted to this stress (Figure 3A). In contrast, no difference was observed in the D-values under N starvation. No such differences were observed when considering the other features. We speculate that these patterns reflect the historical migration of *Prochlorococcus* through the water column over evolutionary timescales. It has previously been suggested that the ancestral LL clades evolved first in deeper waters where nutrient supplies are relatively stable [63]. As the HL clades emerged later to colonize the surface ocean, the severe and dynamic nutrient scarcity there may have acted as a powerful selective pressure, driving the evolution of specialized mechanisms to prolong survival under P-starvation.

**Table 2:** COGs potentially differ in abundance between N and other sensitive strains.

| Proposed process | Annotation | Protein | COG | P value (uncorrected) | More abundant in strains sensitive to | More abundant in strain belonging to |
| --- | --- | --- | --- | --- | --- | --- |
| Quality control-DNA level | Gyrase and DNA topoisomerase | GyrA | COG0188 | 0.012 | N | HL |
|  | Mrr, RecB and HJR activity | Mrr, RecB and HJR | COG1787 | 0.047 | N | - |
|  | very short path repair (Vsr) endocluases | Vsr | COG3727 | 0.024 | N | LL |
|  | DNA ligase I | CDC9 | COG1793 | 0.047 | N | LL |
|  | ArsR family | smtB/ArsR | COG0640 | 0.024 | Other | HL |
|  | type IV methylase | McrA | COG1403 | 0.014 | N | HL |
| Quality control-Protein level | Hsp member- CbpA protein- with DnaJ like domain | CbpA | COG2214 | 0.041 | Other | LL |
|  | Hsp member- Hsp70 | DnaK | COG0443 | 0.034 | Other | LL |
|  | Hsp member- DnaJ chaperone | DnaJ | COG0484 | 0.047 | Other | LL |
|  | Protease- HflB (FtsH) modulator | HflC | COG0330 | 0.041 | Other | LL |
|  | a pseudouridylate synthetase- site specific | RsuA | COG1187 | 0.041 | Other | LL |
| **Amino acid and mixotrophy related biosynthesis** | Amino acids as nitrogen sources- threonine dehydrogenase | EutG | COG1454 | 0.013 | Other | LL |
|  | hydrogenase/ urease accessory protein HupE/UreJ protein | HupE | COG2370 | 0.024 | Other | - |
|  | Asparagine synthetase B | AsnB | COG0367 | 0.028 | N | - |
|  | Chorismate synthase | AroC | COG0082 | 0.041 | Other | LL |
|  | ABC-type bacteriocin/lantibiotic exporters | SunT | COG2274 | 0.042 | Other | LL |
|  | type II secretory pathway | PulO | COG1989 | 0.045 | Other | LL |
|  | membrane-fusion protein | AcrA | COG0845 | 0.042 | Other | LL |
|  | ATPase components of ABC transporters | Uup | COG0488 | 0.047 | Other | LL |
|  | Carbohydrate transport and metabolism, Amino acid transport, and metabolism | RhaT | COG0697 | 0.028 | Other | LL |
|  | ABC transporter involved in maltose transport | MalK | COG3839 | 0.041 | Other | LL |
| **LPS** | CMP-KDO synthetase | kdsB | COG1212 | 0.037 | Other | - |
|  | Carbohydrate transport and metabolism, Amino acid transport, and metabolism | prop | COG0477 | 0.009 | Other | LL |
|  | carbohydrate transport and metabolism | BglX | COG1472 | 0.041 | Other | LL |
|  | Lipopolysaccharide biosynthesis protein | RgpF | COG3754 | 0.038 | Other | LL |
|  | sugar aminotransferase | WecE | COG0399 | 0.048 | N | HL |
|  | poorly characterized, dehydrogenase-sugar aminotransferase fusion protein | MviM | COG0673 | 0.024 | N | - |
| **Membrane linker** | links inner and outer membranes | TonB | COG0810 | 0.041 | Other | LL |
| **Exopolysaccharide** | GumC/Wzc1, integral membrane protein | GumC | COG3206 | 0.009 | N | - |
|  | outer membrane channel, ESP are translocated by it | Wza | COG1596 | 0.004 | N | HL |
| **Biofilm/Attachment** | CsgG protein | CsgG | COG1462 | 0.037 | Other | - |
|  | Large exoprotein involved in heme utilization or adhesion | FhaB | COG3210 | 0.029 | Other | - |
| **Energy production** | Cytochrome c mono- and diheme variants | CccA | COG2010 | 0.048 | Other | - |
| **Translation, transcription, and signal transduction** | Adenylate cyclase | AcyC | COG2114 | 0.038 | Other | - |
|  | Acetyltransferase | PhnO | COG0454 | 0.039 | Other | - |
| **RTX toxins** | RTX toxin-related domain |  | COG2931 | 0.012 | Other | LL |
| **Others** | Dienelactone hydrolase | DLH | COG0412 | 0.013 | Other | - |
|  | Extracytoplasmic sensor domain CHASE2 | CHASE2 | COG4252 | 0.038 | Other | - |
|  | carbamoyl transferase |  | COG2192 | 0.011 | N | HL |
|  | lignin degradation | LigW | COG2159 | 0.002 | N | - |
|  | PPE-repeat proteins | PPE | COG5651 | 0.037 | Other | - |

Because the Benjamini-Hochberg adjustment reaches a mathematical plateau due to our sample sizes and the rank-based nature of the test, the adjusted *P* value is exactly 0.216 across all rows.

**Figure 3:**
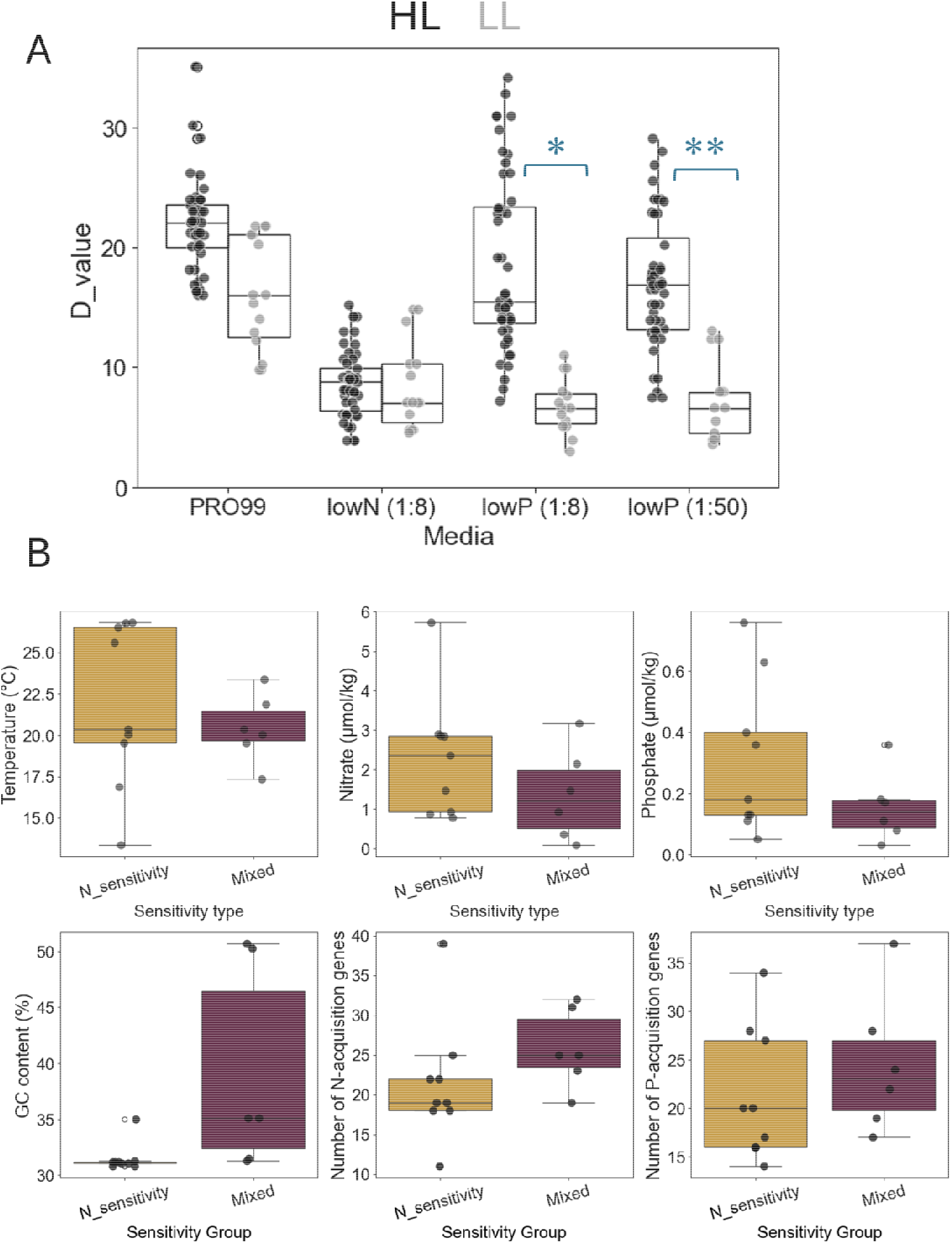
Evolutionary history partially correlates with culture decline whereas ecological history (environmental conditions when and where strains were isolated) do not. A. D-values are higher under P starvation in HL strains; asterisks indicate statistically significant differences. B. Environmental conditions when and where strains were isolated, GC content and number of N and P genes known to be associated with nutrient acquisition in N- and mixed strains, as classified using the D-value. For results based on MaxFL and peak number see Supplementary Table S4.

An alternative hypothesis is that the patterns of mortality represent adaptation of each *Prochlorococcus* strain to the specific local environmental conditions present when and where it was isolated [21]. For instance, the North Atlantic has lower phosphate concentration in surface waters compared to the North Pacific, which is mirrored by the number of genes encoded in the genomes in strains from the region that are associated with N or P uptake [21]. To compare between strains and relate their starvation response to broader environmental and ecological factors, we first divided the strains into two categories, based on their D-value, as it provides a robust and interpretable measure of cell death compared to MaxFL and peak count. Strains that exhibited a higher D-value (slower death rate) under P starvation, survived P starvation better and were consequently classified as N-sensitive whereas those that either died faster under P starvation or equally under N and P were termed as having a mixed sensitivity. Based on that, 9 strains were classified as N-sensitive and 6 strains as mixed. (see phylogenetic tree in Figure 2, Supplementary Table S3, also for classification based on peak count and MaxFL).

Apart from the HL/LL division, the ecological and evolutionary history of the *Prochlorococcus* strains had no clear relationship with the sensitivity to N or mixed sensitivity (Figure 3B). There was no correlation between starvation sensitivity and the environmental conditions (temperature, nitrate, or phosphate concentrations) when and where each strain was isolated, estimated using the World Oceanic Atlas (WOA) ([32], Supplementary Text S5). Moreover, no such correlation was observed when the average seasonal or annual environmental conditions were considered (Supplementary Figure S8, Supplementary Table S5 and Supplementary Text S5). Finally, starvation sensitivity did not correlate with the number of genes associated with N or P uptake (Figure 3B, [20,33–35]). The only correlation observed was with the GC content of the genomes, which reflects the HL/LL division [47]. Taken together, these results suggest that while the deep evolutionary history of the *Prochlorococcus* clade is somehow related to the sensitivity to different types of nutrient starvation, the short-term ecological history of each strain, or the (known) genes for N and P acquisition encoded in its genome, are unlikely to drive the distinct death patterns observed (Figure 3A, Supplementary Table S4).

### Identifying genes that are potentially associated with starvation sensitivity

We next searched for genes that are differentially distributed between N and mixed strains, and that could hint at the molecular mechanisms underlying the differential mortality patterns. To do so, we examined the abundance of genes belonging to different Clusters of Orthologous Genes (COGs) across the *Prochlorococcus* genomes studied, using CyanoRak [40]. There are 1,380 COGs in the *Prochlorococcus* pan genome, of which 670 varied in gene number between strains. 112 of these differed in number between HL/LL strains or N- and mixed sensitivity ones (Figure 4A).

**Figure 4:**
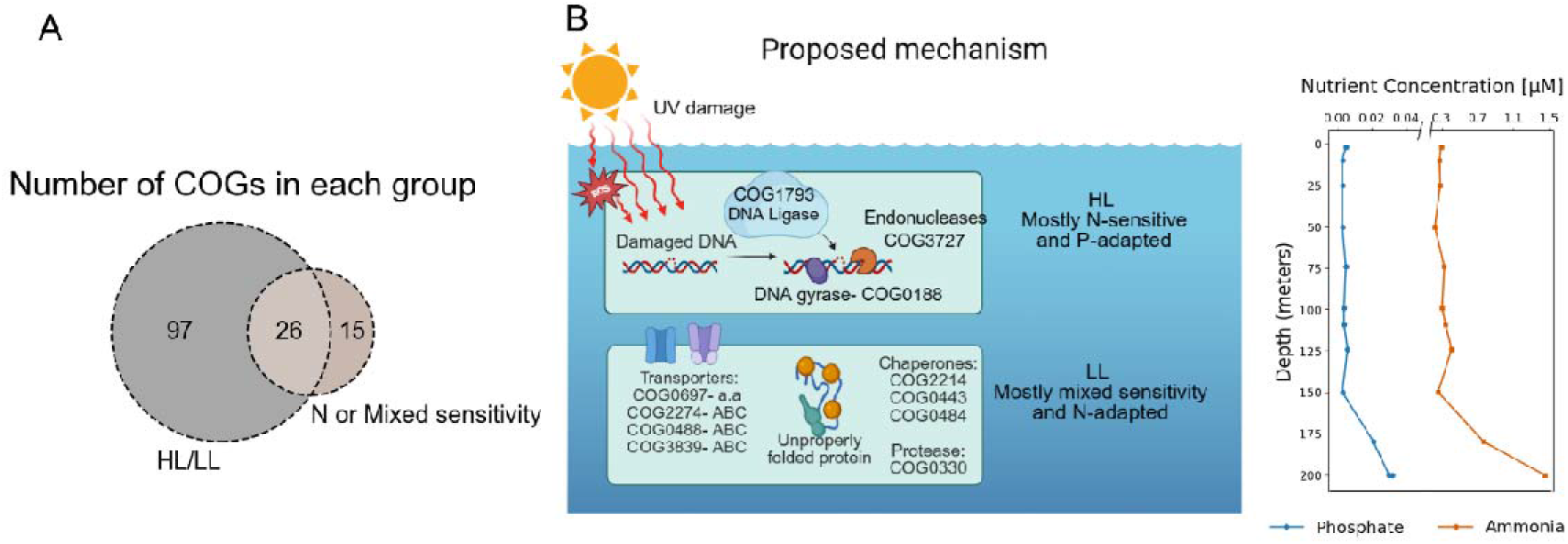
Proposed cellular mechanisms underlying starvation sensitivity in *Prochlorococcus* and their division between HL and LL clades. A. The number of COGs differing between HL and LL strains and/or between N and mixed-sensitive ones (*P* <0.05, uncorrected for multiple hypothesis testing). B. An illustration showing proposed key cellular mechanisms underlying death in *Prochlorococcus* on the left, and a nutrient profile for phosphate and ammonia from a cruise in the Eastern Mediterranean on the right, demonstrating nutrient concentrations in regions inhabited by HL and LL strains.

Although some COGs appeared to be associated with N or mixed sensitivity, the differences in gene count within each COG were relatively small. In fact, no significant associations remained after applying for the Mann-Whitney U test followed by the Benjamini-Hochberg correction for multiple testing. Additionally, interpretation of the results was complicated by the cross-correlation between P sensitivity and LL strains (Figure 3A). Despite this, certain patterns emerged among the COGs linked to specific types of starvation sensitivity. These patterns may offer insight into potential mechanisms underlying the mortality responses observed. Therefore, we highlight the COGs that showed significance at *P* < 0.05 without correction for multiple testing (Table 2 and Dataset 3).

Overall, before multiple hypothesis correction, 26 COGs were associated, with both starvation type and evolutionary history (the HL/LL division), and another 15 were only associated with starvation sensitivity (Figure 4A, Table 2). The pattern that was most clearly associated with sensitivity to N starvation was an increased number of genes involved in DNA damage control, namely DNA gyrase (COG0188), helicase/nuclease (*recB*, COG1787), endonucleases (COG3727), and DNA ligase (COG1793), (Table 2, Figure 4B). The correlation offers two potential interpretations. First, it is possible that a higher copy number of these DNA repair genes directly correlates with increased sensitivity to N starvation. Alternatively, because sensitivity to N and P starvation are inversely related in our dataset, this genomic enrichment may instead reflect a mechanism that allows strains to better cope with P starvation. Under this alternative view, having more copies of these genes is an asset during P stress, rather than a liability during N stress. In support of the second interpretation, the enrichment of DNA damage repair genes in HL strains aligns with their ecological niche, as these strains predominantly inhabit the ocean’s surface, where nutrient availability is limited and exposure to UV radiation and reactive oxygen species (ROS) is high. These environmental stressors are strongly linked to DNA damage [66,67], necessitating an enhanced capacity for DNA repair. Further support is provided by the observation that two HL strains, SB and MIT9301, which were mixed sensitive rather than N-sensitive, had lower numbers of the COGs associated with DNA damage compared with other HL strains.

In contrast, strains more sensitive to mixed starvation exhibited a greater abundance of genes associated with protein quality control including chaperones (COG2214, COG0443, and COG0484) and a protease (COG0330), (Table 2, Figure 4B). These genes were also more abundant in LL strains. The enrichment of these protein quality control systems may reflect an enhanced capacity to manage protein stress during nutrient limitation. Chaperones and proteases play a documented role in mitigating protein misfolding, aggregation, and degradation under stress conditions [68]. which might contribute to the observed differences in survival under N starvation. However, the specific physiological advantage this provides, and why these functions are also more abundant in LL strains, requires further investigation.

Genes related to mixotrophy and/or alternative nitrogen sources, such as amino acid metabolism (COG1454) and a urease subunit (COG2370), were more abundant in mixed sensitive strains, but also in LL strains. As noted above, in principle a higher number of such genes could suggest better resistance to N starvation through mixotrophy [60], LL strains and HL strains die at the same rate under N starvation (Figure 3A), and thus increased mixotrophy does not seem to be a strong adaptive strategy to cope with this stress. Perhaps this is due to the experimental conditions, where the closed system lacks a continuous flux of organic nutrients to serve as a substrate. Alternatively, under sudden, acute starvation, cells may become rapidly depleted of the ATP required to power active, energy-dependent organic nutrient transporters, rendering them unable to utilize these alternative reservoirs, some of which likely become available as cells die and lyse. Genes related to lipopolysaccharides, exopolysaccharides, biofilm attachment, energy production, translation, transcription, and signal transduction were also differentially associated with either N or mixed sensitivity, but these associations were less consistent (Table 2), and we currently cannot propose an explanation for them.

Unlike the known N/P acquisition genes, which were differentially expressed (as a group) in cultures starved for up to 48-59 hours (for N and P, respectively), the COGs associated with N or mixed sensitivity were not clearly differentially expressed under these starvation conditions (Supplementary Text S6, Supplementary Figure S9). Instead, many of the COGs, especially those associated with DNA quality control, were more abundantly expressed under extended N starvation (4 days after starvation began, [70]) This suggests that these genes may be associated with later stages of starvation as the cells begin to die.

## Conclusion, caveats, and prospects

Death due to nutrient starvation remains poorly understood in microbes in general, and in marine bacteria (including cyanobacteria) in particular. In this study, we investigated features of starvation and mortality-maximum fluorescence, number of peaks, and death rates-across diverse *Prochlorococcus* strains under N and P starvation. These measurements revealed complex, strain-specific patterns of starvation survival. Although lowP (1:50) consistently resulted in the lowest biomass yield, this did not necessarily correlate with the shortest survival time; in contrast, N-deplete conditions induced a sharp, rapid decline across all strains. These diverging responses suggest that N-limitation imposes a more immediate mortality constraint compared to P-limitation, raising new hypotheses regarding the distinct molecular programs that govern starvation resilience in these microbes.

The patterns of mortality due to starvation were influenced by the strains’ evolutionary history (HL/LL divide), but not, as far as we could resolve with the culture collection in our hand, by their shorter-term ecological history (based on the conditions where they were isolated from or the number of previously characterized genes involved in N and P acquisition encoded in their genomes). This finding stands in contrast to the expectation that N and P acquisition genes have been considered as markers for microbial survival and adaptivity in nutrient-limited environments [20,22,34]. Furthermore, N acquisition genes have been linked to phylogeny, whereas P acquisition genes exhibit a correlation with environmental location [23,69]. Mapping the evolutionary and ecological history of the strains was challenging, as the record of when and where many strains were isolated, and what the environmental conditions were at the time, was partial or missing. Systematically collecting and reporting such data is necessary for future work to integrate phenotypic, genomic, and environmental data.

Even though our study focuses on overall growth and mortality, we did not directly measure the real-time physiological response to starvation. Bacteria often rely on the stringent response, a survival mechanism that slows down cellular processes like protein production and DNA replication, to endure nutrient-limited conditions [71]. Two of the genes we identified (COG0188/DNA gyrase and COG1787/RecB helicase) are known to help regulate this response in other model organisms by inhibiting DNA replication [72]. It is possible that variations in this starvation-survival program contribute to the differences in mortality we observed between strains. Future research should track how these molecular pathways shift as cells transition from survival to death, which will be essential for understanding the life-and-death strategies of microbes in nutrient-poor ocean environments.

The genomic copy number of mixotrophy-related genes did not influence mortality rates. This was contrary to our expectation that a broader capacity to utilize organic substrates would provide LL strains with a clear survival advantage during nitrogen and phosphorus starvation (Table 3, [60,61]) but is consistent with mathematical modeling suggesting that mixotrophy does not underly slower mortality in co-cultures with heterotrophic bacteria [27]. Instead, we identified genes related to protein and DNA quality control that might be associated with the rate of *Prochlorococcus* mortality in response to N or P starvation. These associations suggest that protein homeostasis may play a key role in LL strains, whereas DNA repair mechanisms are particularly important for managing P-starvation, primarily in HL strains. Currently, there are no methods for gene manipulation in *Prochlorococcus* [73], and therefore testing these hypotheses will either have to wait for such methods to be developed, or they will need to be tested in related cyanobacteria such as *Synechococcus*.

## Data and code availability

Because no public repository currently maintains these axenic strains, they are available from the authors upon request.

All the code and data used in this manuscript is available on https://github.com/TheSherLab/15strain4media_analysis

## Acknowledgments

We would like to thank the Chisholm lab team for supplying the cultures used in this study and for insightful discussions, and Daniel Vaulot, Frederic Partensky and Laurence Garczarek for assistance in identifying the environmental conditions when strains were isolated, and for providing unpublished data from Cyanorak.

This work was funded by the Israel Science Foundation (grant number 1786/20 to DS) and by the National Science Foundation - United States-Israel Binational Science Foundation (NSFOCE-BSF 1635070 and NSF-BSF 2246707 to DS). Large language models were used for language editing, to assist with modifying some of the code related to figure generation, and for the integrated transcriptome analysis. The funders had no role in study design, data collection and analysis, decision to publish, or preparation of the manuscript.

