## Supplementary text, tables and figures for "Dynamics of cell death due to N and P starvation across intra-clade diversity in *Prochlorococcus*"

Yara Soussan-Farhat, Shira Givati, Osnat Weissberg, Waseem Bashir Valiya Kalladi, Dikla Aharonovich and Daniel Sher
Department of Marine Biology, Leon H. Charney School of Marine Sciences, University of Haifa, Haifa, Israel.

**Supplementary Text S1: Fluorescence (FL) as a proxy for cell abundance.**

Growth-and-death curves are biologically complex. Although bulk fluorescence (FL) is a good determination of growth in balanced exponential growth, one needs to assess the relationship between FL and cell count under N and P limitation. For that, we performed two separate experiments:

1. The "cell abundance experiment". In this experiment, four strains (MIT9312, MED4, SB, and MIT1314) were grown across the four media (described in Supplementary Table S1), their FL was measured across 99 days, and samples for cell abundance were taken at different stages of the curve, mainly at the decline stage.
2. The "spiking experiment" (discussed in detail in Supplementary Text S2), which included five strains (MIT9312, MED4, SB, MIT1314, and PAC1) and all media, but where the sampling focused on exponential growth and the peak.

In both experiments cultures were grown under constant light (22-25 µmole photons $m^{-2}$ $s^{-1}$) at 22°C. Combined, this resulted in a total of $\sim 310$ sampling points checked for cell abundance by flow cytometry (see Material and methods).

A linear relation was observed between cell abundance, measured as cells $mL^{-1}$, and bulk FL (Supplementary Figure S1A ), with an F-statistics= 641.7, $R^2$ =0.67, and $P$ value of $6.75\times10^{-77}$, consistent with previous studies [1]. No difference was observed in the slope of the regression between media types, suggesting that growth and death parameters are comparable between nutrient types, at least for individual strains. However, the slope is different between some strains. This means that even though cell numbers can be calculated to some extent from FL measurements from the same strain, differences between strains in the slope of the regression make such comparisons difficult across strains (Supplementary Figure S1B).

Despite the overall correlation between cell numbers and bulk culture fluorescence, previous studies have shown that per-cell fluorescence decreases as cultures stop growing and start to decline (chlorosis,[2]). Because we would like to differentiate, if possible, between changes in cell numbers and chlorosis at this stage, we zoomed in on the points that only represented the peak from the "spiking experiment" (Supplementary Figure S1C). We see that the linear relation between cell numbers and culture fluorescence is still present, and the difference between strains is reflected in the slopes. It also shows that cell abundance is the major player driving the differences in $Max_{FL}$ within each strain. The FL per cell is maintained at the peak, whereas the absolute cell abundance is different. One strain, PAC1, shows a different behaviour were FL per cell changes across media. We believe this is largely due to the elongated scatter of the culture and to it being a low-light (LL) strain, which is characterized by larger cells (compared to the high-light strains) and therefore has more chlorophyll within each cell. This shows that the differences we see in $Max_{FL}$ are mirrored by similar differences in cell numbers across both N and P limitations.

What about the decline stage? Is the drop in bulk FL mainly due to cell death or due to chlorosis, which is accompanied by a decrease of FL per cell? Because we do not discount that chlorosis plays a part during decline, we tested the actual changes in bulk FL, cell abundance, and FL per cell within this stage. We chose two points in the decline phase and calculated the fold change by dividing the earlier value by the later value for all three parameters to test the relation between them. shows that Bulk FL fold decline is linearly correlated with the cell abundance fold change, but not with the FL per cell fold change (Supplementary Figure S1D). Even though this does not mean chlorosis is entirely absent, it provides direct evidence that cell death is the primary driver of the bulk fluorescence decline observed in our curves, justifying the use of bulk FL for calculating the D-value.

Lastly, we do not have sufficient data to systematically test whether the multiple peaks observed (primarily in the PRO99 media) correspond to changes in cell number or FL per cell. Nevertheless, we have observed several cases where multiple peaks were clearly present in the absolute cell numbers as well (Supplementary Figure S1F). Therefore, in at least some cases, these recurring fluorescence peaks represent true population re-growth rather than optical or physiological artifacts.

**Supplementary Text S2: Spiking cultures with limiting nutrients.**

An experiment was set up using five strains under four growth conditions (this experiment is referred to as the spiking experiment): PRO99, lowN (1:8), lowP (1:8), and lowP (1:50). Four replicates were included for each strain/medium combination. Cultures were grown under constant light (22-25 µmole photons $m^{-2}$ $s^{-1}$) at 22°C. When a strain/medium combination showed signs of decline, two replicates were spiked with the limiting nutrient to match PRO99 concentrations, whereas the other two replicates were left untouched.

The spiked cultures ceased their decline and resumed growth (Supplementary Figure S2). The $Max_{FL}$ of the spiked cultures also increased to match that of PRO99, confirming that $Max_{FL}$ is directly affected by the nutrient starvation the culture is experiencing (Supplementary Figure S2B). PRO99 control tubes were also spiked with both N and P, but no additional growth was observed, showing that the decline under PRO99 was not due to nutrient limitation.

All strain under lowP (1:8) failed to show subsequent growth after the addition of P, emphasizing that they were not completely starved by the 8-fold P reduction. PAC1 under lowP (1:50) also failed to resume growth. We speculate this is due to extreme starvation, where the cultures reached a "point of no return" and could not recover from the severe stress. This aligns with observations from other experiments where transferring cultures into fresh, nutrient-rich media after 3-4 days of decline did not result in regrowth [2].

**Supplementary Text S3: Comparison between the spiking experiment and the main experiment, and their axenicity test.**

Since the initial experiment only included PROMM to check for axenicity, we decided to expand this in the spiking experiment. Here, we used both PROMM and MB media at the start and end of the experiment as indicators for contamination; tubes were incubated under identical conditions to the experimental setup, and cultures were visually examined for turbidity after 7 days. Any strain exhibiting visible growth was classified as non-axenic and excluded from the analysis.

Additionally, we sequenced the 16S rRNA genes (V4-V5 region, primers 515F-Y [5′-GTGYCAGCMGCCGCGGTAA-3′] and 926R [5′-CCGYCAATTYMTTTRAGTT-3′]) on a MiSeq System (Ilumina) to obtain a community profile. Axenic *Prochlorococcus* cultures are expected to yield a single dominant ASV, with very low levels of other "contaminating" ASVs resulting from sequencing errors or cross-contamination during DNA extraction, amplification, or sequencing steps. We operationally defined a true contaminant as a non-*Prochlorococcus* ASV with a relative abundance of >0.3%. In all samples, the maximum abundance of the second most abundant ASV was 1.5%; these were mostly identified as *Prochlorococcus* and are thus likely sequencing errors. In contrast, non-axenic positive control cultures had 10.6–19.4% of reads originating from other organisms.

Because the spiking experiments resulted in patterns similar to the initial experiment (Supplementary Figure S3) and confirmed axenicity, we conclude that the patterns observed in the original experiments were likewise unaffected by contamination.

**Supplementary Text S4: Growth curves of MED4 and MIT9312 in four independent experiments.**

Due to the large number of strains and conditions tested, we performed four different experiments, each with different strains. In order to make sure that there is no “batch effect” (i.e. that observed differences in starvation and mortality between strains are not in fact due to experiment-to-experiment variation), we included in each experiment two strains, MED4 and MIT9312, as internal standards. In some experiments a short lag phase was observed, for example under lowN (1:8) and lowP (1:50) in experiment 1 for MED4. This could be due to slightly different timing of transfer from phase 1 (acclimation to low nutrient media) to phase 2 (the actual experiment). However, the death phase, which was the primary focus of this study, exhibited consistent patterns and high reproducibility (Figure 2, Supplementary Figure S4).

**Supplementary Text S5: The environmental measurements tested here were monthly interpolated data from the World Oceanic Atlas (WOA).**

Because complete environmental data were not available for all strains from the cruises on which they were isolated, we supplemented our dataset with information from WOA. To ensure consistency, we first compared the available strain-specific data with the corresponding WOA data (Supplementary Figure S7A) and found no significant variability.

The WOA provides data at monthly, seasonal, and annual resolutions. To assess the impact of these different timescales, we conducted comparative analyses. Nitrate and phosphate measurements exhibited some differences (Supplementary Figure S7B). However, no significant correlations were observed when mean seasonal and annual conditions were used (Supplementary Table S4). Based on these findings, we selected monthly data for our analysis, as it provides a representative and reliable resolution for our study.

**Supplementary Text S6: Gene- and COG-level expression response to N and P starvation experiments in vivo, using a knowledge graph (KG).**

N and P acquisition genes were thought to be essential for coping with N and P starvation, respectively. In our study we showed that the copy number of these genes per strain did not reflect the death patterns observed. To assess whether these genes actually respond to nutrient starvation in vivo, we used a knowledge-graph-based system (see Material and methods), to test whether these genes were up or downregulated in multiple N and P starvation experiments. In total we found 10 individual starvation experiments, five for N and five for P, reported in six studies [2-8]. With these data we were able to test 60 of the 92 previously selected N or P responsive genes (these 60 can be related to a specific locus tag in the four strains covered by the 10 experiments (MED4, MIT9312, MIT9313, and NATL2A).

As expected, N acquisition genes were mostly upregulated under N starvation, and P acquisition genes were upregulated under P starvation. We also observed that N genes were not differentially expressed under P starvation, and vice versa. These data reflected experiments where cultures were axenic and used a single representative starvation timepoint for each condition (Supplementary Figure S9A). Furthermore, we tested whether N and P acquisition genes were up- or downregulated compared to randomly selected genes. Across 10,000 iterations including ~100 randomly chosen genes, the N and P acquisition genes fell outside this distribution.

We also tested genes belonging to the COGs associated with starvation sensitivity using the same approach as the N/P acquisition genes. Of the 41 COGs, 37 were represented at least once in the dataset, while the remaining 4 (AsnB, EutG, GumC, Vsr) are found in strains not studied in the transcriptome experiments. In contrast to the N and P acquisition genes, the COGs associated with N and mixed sensitivity were not statistically enriched (up- or down-regulated as a group) under N or P starvation, compared to randomly selected genes. However, DNA quality control genes are mostly upregulated under long term N starvation (4 days, [3]). This requires further investigations.

**Supplementary tables:**

**Supplementary Table S1**: **Nutrient concentrations and their ratios in the different media.**

| **Media** | **Ammonium-N (µM)** | **Phosphate- P (µM)** | **N/P ratio** |
|---|---|---|---|
| **PRO99** | 800 | 50 | 16 |
| **lowN (1:8)** | 100 | 50 | 2 |
| **lowP (1:8)** | 800 | 6.25 | 128 |
| **lowP (1:50)** | 800 | 1 | 800 |

**Supplementary Table S2: Ordinary least squares (OLS) statistical test with multi test correction using the Benjamini–Hochberg method results of death features across the tested strains**.

| | | | | | | | | | | | | |
|---|---|---|---|---|---|---|---|---|---|---|---|---|
| **MIT1327** | a | b | c | d | NS | NS | NS | NS | a | b | b | b |
| **MIT9313** | a | b | b | c | a | b | b | b | a | b | b | b |
| **PAC1** | a | b | a | c | a | b | b | b | a | a | b | b |
| **NATL2A** | a | b | c | d | a | b | a | b | a | b | c | c |
| **NATL1A** | a | b | ab | c | NS | NS | NS | NS | a | b | c | d |
| **MIT1314** | a | b | c | d | a | b | b | b | a | b | c | c |
| **MIT9312** | a | b | b | c | a | b | b | b | a | b | c | c |
| **AS9601** | a | b | a | c | a | b | c | bc | a | b | a | c |

| Strain/Medium | Feature | MED4 | MIT9515 | MIT9202 | MIT9215 | MIT0604 | MIT9301 | SB |
|---|---|---|---|---|---|---|---|---|
| PRO99 | $Max_{FL}$ | a | a | a | a | a | a | a |
| lowN | | b | b | b | b | b | b | b |
| lowP (1:8) | | a | c | a | b | a | c | c |
| lowP (1:50) | | c | d | c | c | c | d | d |
| PRO99 | Peak count | a | a | a | a | NS | a | a |
| lowN | | b | b | b | b | NS | b | ab |
| lowP (1:8) | | a | b | ac | ab | NS | b | b |
| lowP (1:50) | | c | b | bc | b | NS | b | b |
| PRO99 | D-value | a | a | a | a | a | a | a |
| lowN | | b | b | b | b | b | b | b |
| lowP (1:8) | | c | c | c | c | c | b | c |
| lowP (1:50) | | c | c | d | c | b | b | d |

**Supplementary Table S3: Sensitivity of a strain according to each one of the features.**

| Strain/feature | $Max_{FL}$ | Peak count | D-value |
|---|---|---|---|
| **MED4** | N | N | N |
| **MIT9515** | N | N/P | N |
| **MIT9202** | N | N | N |
| **MIT9215** | N/P | N | N |
| **MIT0604** | N | N/S | N |
| **MIT9301** | N/P | N/P | N/P |

| | | | |
|---|---|---|---|
| **SB** | P | P | P |
| **AS9601** | N | N | N |
| **MIT9312** | N/P | N/P | N |
| **MIT1314** | P | N/P | N |
| **NATL1A** | N/P | N/S | N |
| **NATL2A** | N | N | P |
| **PAC1** | N | N/P | P |
| **MIT9313** | N/P | N/P | N/P |
| **MIT1327** | N | N/S | N/P |

**Supplementary Table S4: Statistical tests showing the correlation between the N vs. Mixed sensitive groups and the measurements included.**

| | ***P* value results when testing N vs. Other sensitive according to** | | |
|---|---|---|---|
| **Measurement** | **$Max_{FL}$** | **Peak count** | **D-value** |
| **Temperature** | 0.49 | 0.43 | 0.68 |
| **Nitrate** | 0.56 | 0.72 | 0.32 |
| **Phosphate** | 0.68 | 0.07 | 0.24 |
| **GC%** | 0.73 | 0.38 | 0.01 |
| **N acquisition genes** | 1.00 | 0.72 | 0.08 |
| **P acquisition genes** | 0.20 | 0.35 | 0.31 |

**Supplementary Table S5: Statistical tests showing the correlation between the D-values and the interpolated environmental measurements from the World Oceanic Atlas (WOA).**

| | ***P* value results when testing D-value against environmental measurements** | | |
|---|---|---|---|
| **Measurement** | **Monthly** | **Seasonal** | **Annual** |
| **Temperature** | 0.68 | 0.52 | 0.60 |
| **Nitrate** | 0.32 | 0.28 | 0.32 |
| **Phosphate** | 0.24 | 0.26 | 0.39 |

**Supplementary figures:**

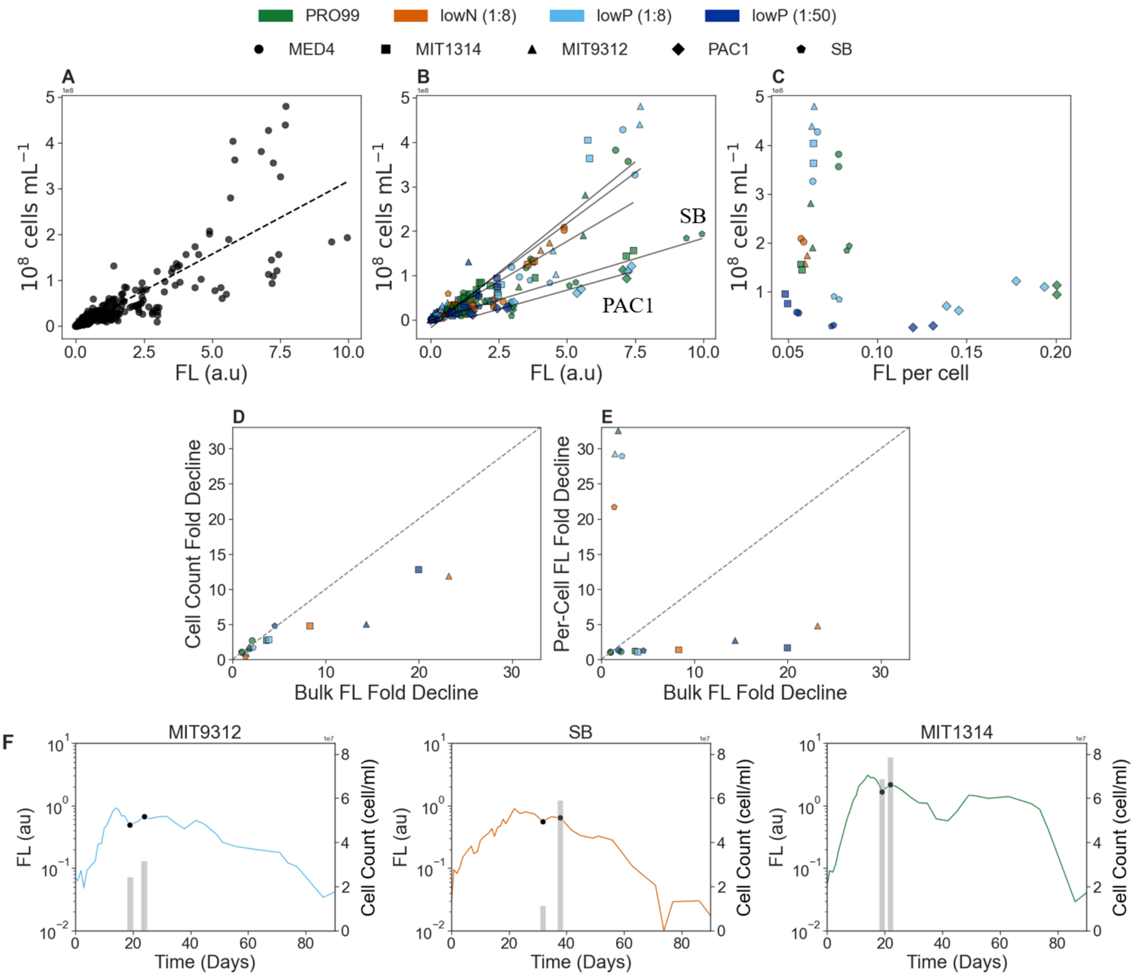


Supplementary Figure S1. Relationship between bulk fluorescence, cell abundance, and per-cell fluorescence across diverse *Prochlorococcus* strains. A. Linear relationship between cell abundance and bulk fluorescence (FL) across all samples, showing a consistent global trend. B. The effects of nutrient conditions on the FL-cell abundance relationship; although nutrient type does not significantly alter the linearity, distinct *Prochlorococcus* strains exhibit variation in their respective regression slopes. C. Subset analysis of the peak phase in spiking experiments, demonstrating that differences in bulk FL across nutrient conditions are primarily driven by variations in cell abundance. D, E. The relations between fold-change in cell abundance (D) and per-cell fluorescence (E) as a function of bulk FL decline. F. Representative examples highlighting instances during the decline phase where subsequent peaks showing coupled increase in bulk FL and cell abundance.

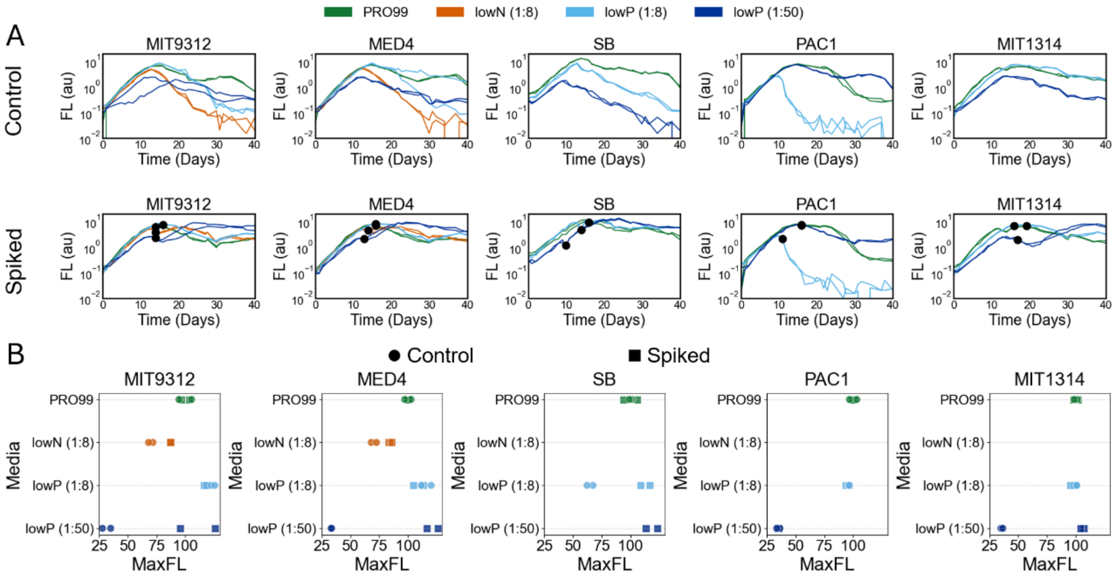


Supplementary Figure S2: Growth-and-death curves of five selected strains. A. The x-axis represents time in days, and the y-axis shows fluorescence (FL), a proxy for cell biomass, on a logarithmic scale. Colors indicate the different nutrient media. The upper row displays two biological replicates that were left un-spiked, the lower row shows replicates spiked with the limiting nutrient. The black dots in the lower row mark the days the limiting nutrient was added. B. Changes in $Max_{FL}$ between control and spiked replicates.
The lack of lowN data for SB, PAC1, and MIT1314 is due to the slower initial growth of these strains under lowN at the time the experiment was conducted.

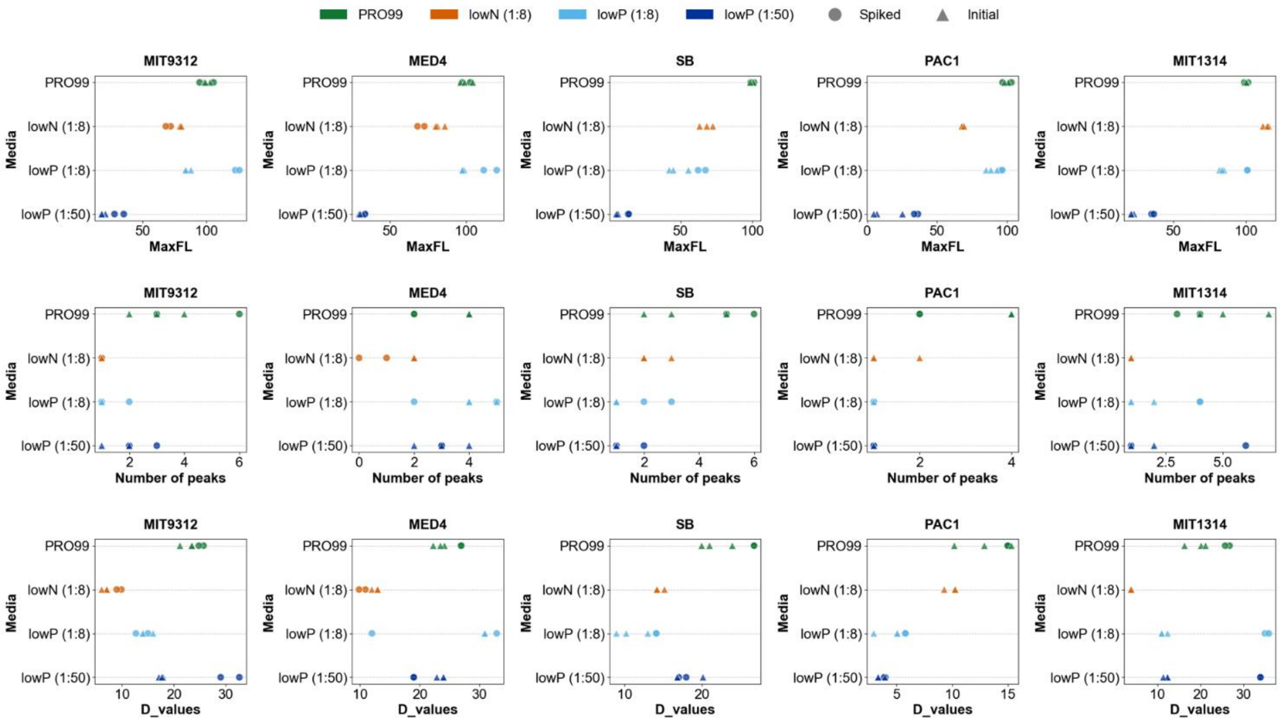


Supplementary Figure S3: Comparison between the initial and spiking experiments. Different rows show the results of three features ($Max_{FL}$, number of peaks, and D- value) calculated in both experiments, demonstrating that the observed patterns are largely similar and that the two experiments are comparable.

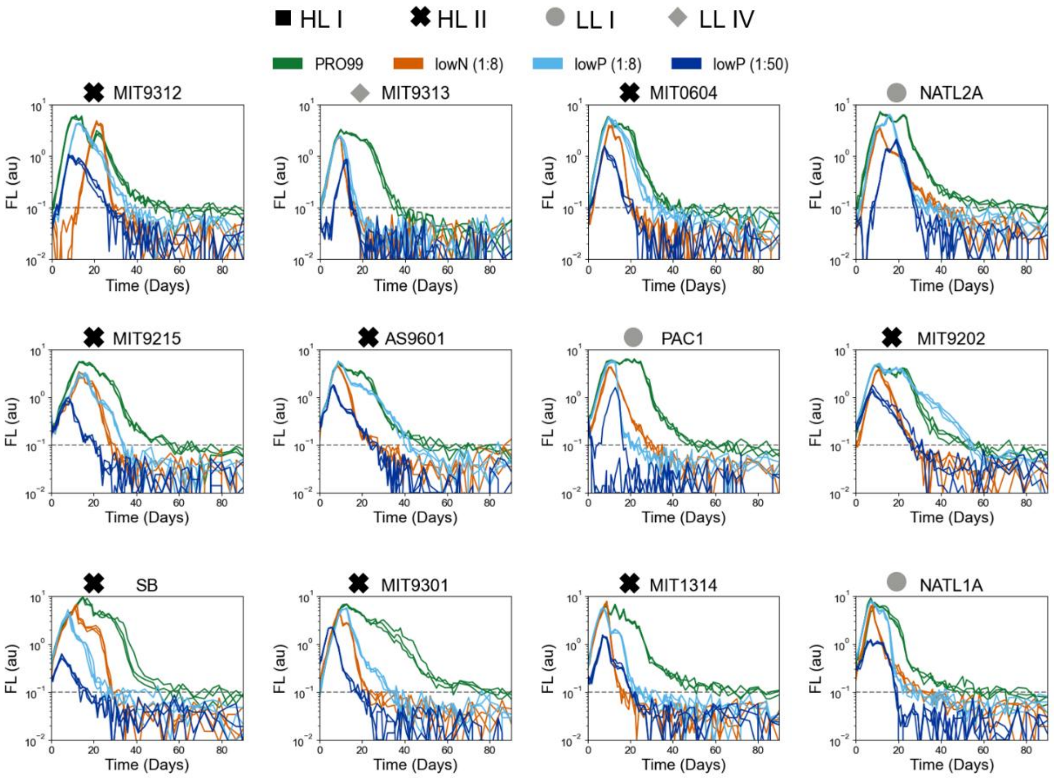


Supplementary Figure S4: Growth curves of 12 strains, the x-axis represents time in days and the y-axis is FL, a proxy for cell growth, on a logarithmic scale. Colors represent the nutrient media cultures grow under, whereas shapes describe the clades. The curve of MIT9312 is from a representative individual experiment.

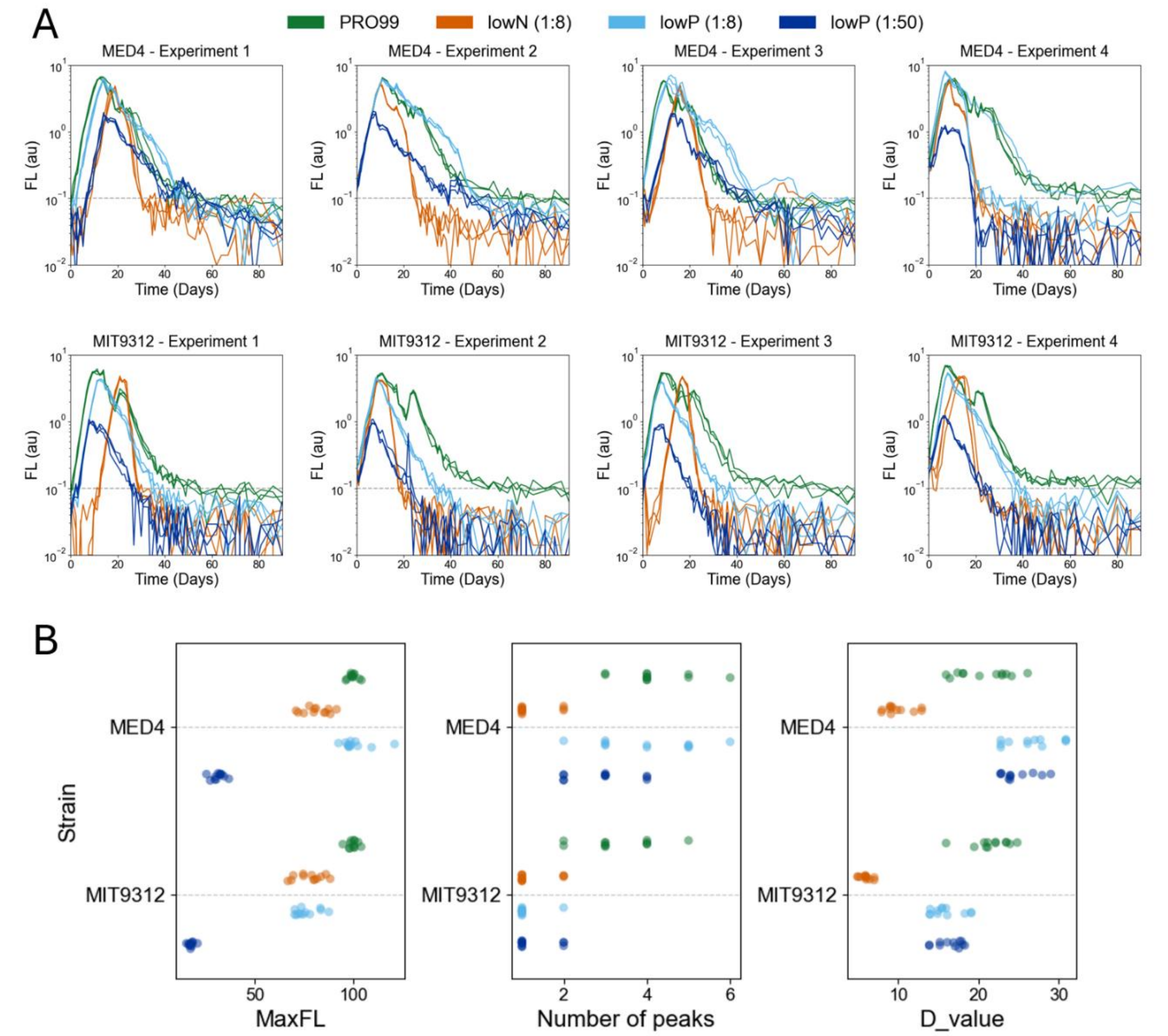


Supplementary Figure S5: Growth-and-death curves of MED4 and MIT9312 in four independent experiments. A. Growth curves of MED4 and MIT9312 in the four independent experiments. B. Measurements of the three features; $Max_{FL}$, number of peaks and D-value across all four experiments in MED4 and MIT9312.

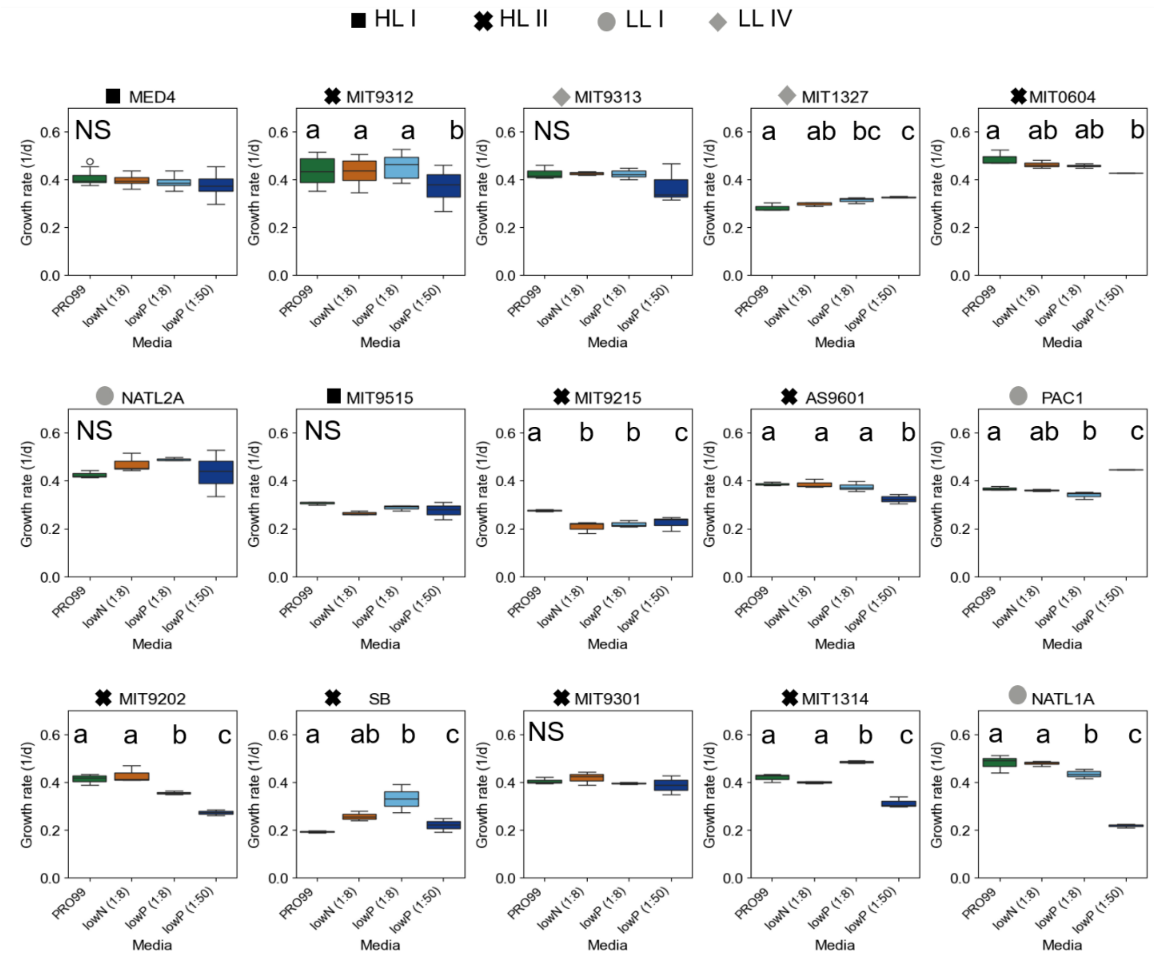


Supplementary Figure S6: Growth rates result across media in each strain. Colors represent the different media. Statistical test results are overlayed. Not significant (N/S) marks strains with no differences across media.

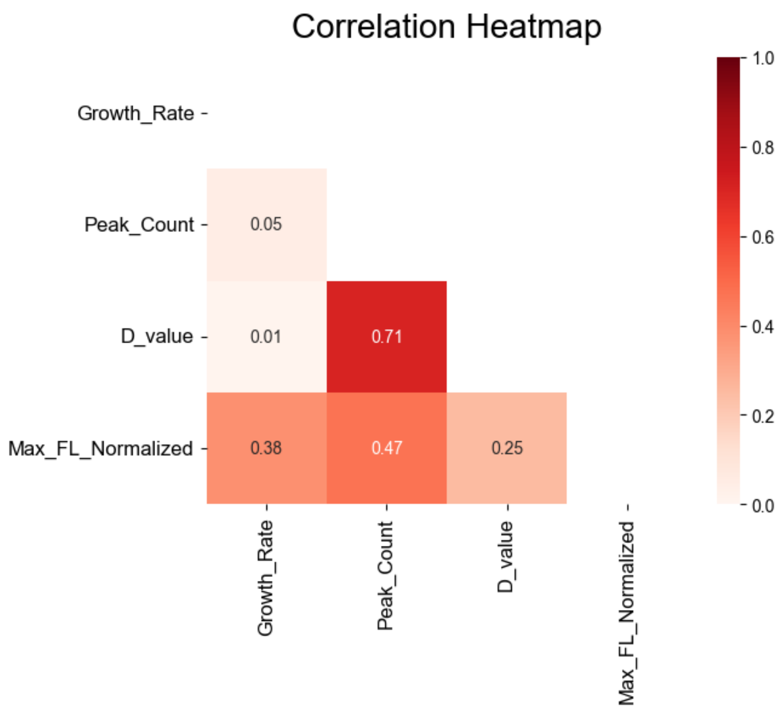


Supplementary Figure S7: Pearson correlations coefficient between the distinct death features.

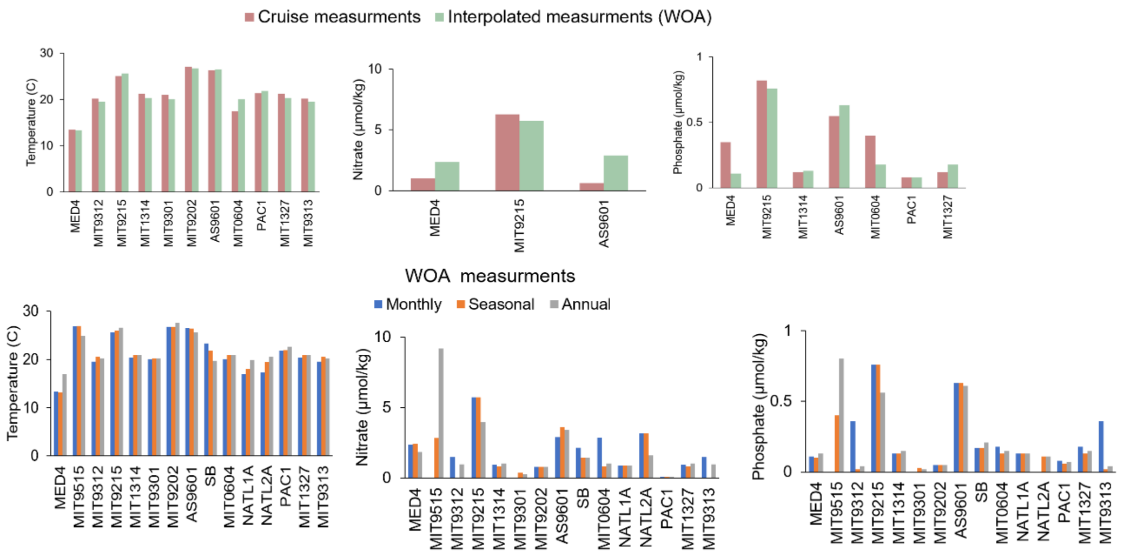


Supplementary Figure S8: Compresence between data from cruises and interpolated data from the World Oceanic Atlas (WOA).

A. Cruise data vs. interpolated data. B. Comparisons between monthly, seasonal, and annual data from the WOA.

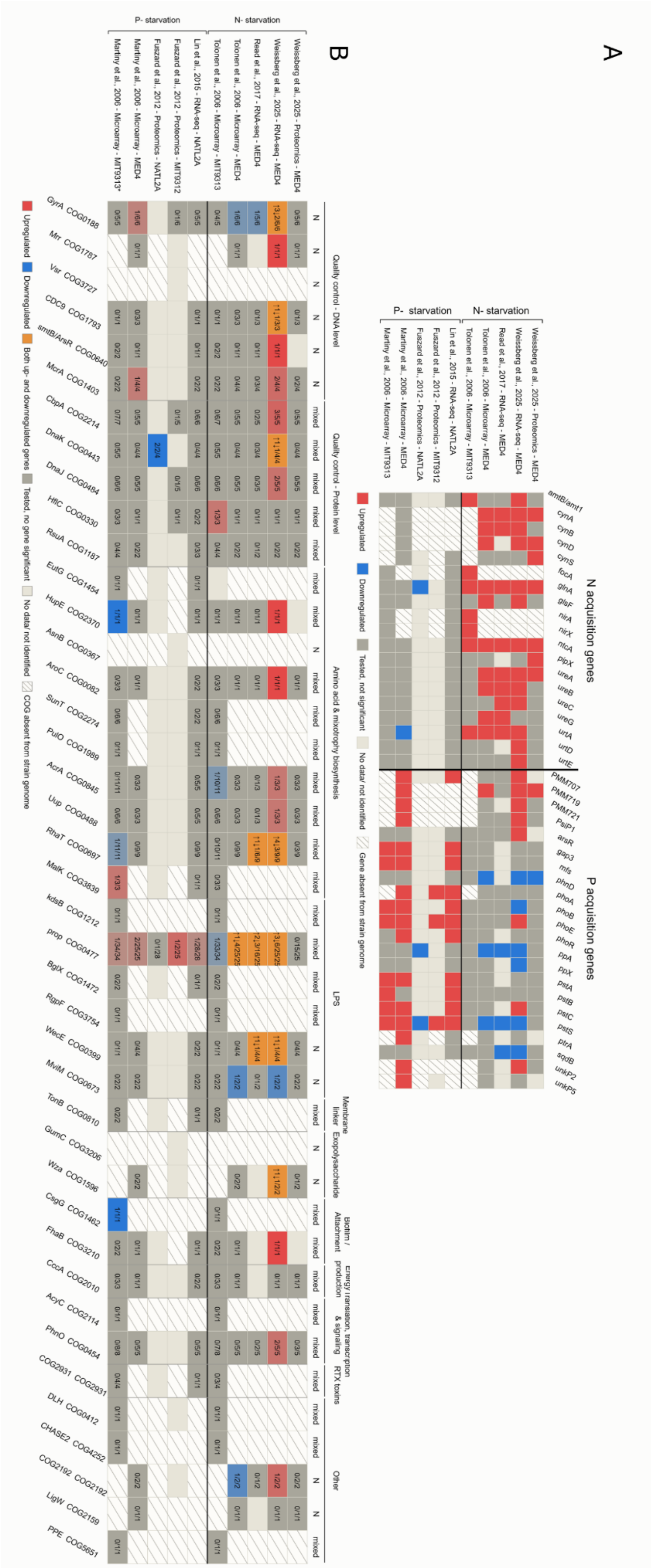


Supplementary Figure S9: In vivo expression response to nutrient starvation: N/P-acquisition genes (A) and starvation-sensitivity-associated COGs (B) in *Prochlorococcus.* The data were collected

into a Knowledge Graph (KG), and the fold change and significance extracted from the initial publications. The three numbers shown in panel B represent number of genes or proteins significantly differentially expressed in the analysis/number of genes tested in the analysis/number of genes in the specific COG. If part of the genes in the same COG was up and others where down regulated, the cell is marked with orange. The number of upregulated genes are shown with and up arrow and these that are down regulated with a down arrow.
Proteomic analysis identifies only a subset of a strain's proteins; therefore, many genes simply have no protein-level data at all in Weissberg et al., 2025 and Fuszard et al., 2012.
P starvation studies report data for the significantly differentially expressed genes only, set as q < 0.05 in Martiny et al., 2006 and *P* adj < 0.05 with |log2FC| > in Lin et al., 2016.
